# One dose of COVID-19 nanoparticle vaccine REVC-128 provides protection against SARS-CoV-2 challenge at two weeks post immunization

**DOI:** 10.1101/2021.04.02.438218

**Authors:** Maggie Gu, Jonathan L. Torres, Jack Greenhouse, Shannon Wallace, Chi-I Chiang, Abigail M. Jackson, Maciel Porto, Swagata Kar, Yuxing Li, Andrew B. Ward, Yimeng Wang

## Abstract

A COVID-19 vaccine with capability to induce early protection is needed to efficiently eliminate viral spread. Here, we demonstrate the development of a nanoparticle vaccine candidate, REVC-128, in which multiple trimeric spike ectodomain subunits with glycine (G) at position 614 were multimerized onto a nanoparticle. In-vitro characterization of this vaccine confirms its structural and antigenic integrity. In-vivo immunogenicity evaluation in mice indicates that a single dose of this vaccine induces potent serum neutralizing antibody titer at two weeks post immunization, which is significantly higher than titer induced by trimeric spike protein without nanoparticle presentation. The comparison of serum binding to spike subunits between animals immunized by spike with and without nanoparticle presentation indicates that nanoparticle prefers the display of spike RBD (Receptor-Binding Domain) over S2 subunit, likely resulting in a more neutralizing but less cross-reactive antibody response. Moreover, a Syrian golden hamster in-vivo model for SARS-CoV-2 virus challenge was implemented at two weeks post a single dose of REVC-128 immunization. The results show that vaccination protects hamsters against SARS-CoV-2 virus challenge with evidence of steady body weight, suppressed viral loads and alleviation of tissue damage (lung and nares) for protected animals, compared with ~10% weight loss, higher viral loads and tissue damage in unprotected animals. Furthermore, the data show that vaccine REVC-128 is thermostable at up to 37°C for at least 4 weeks. These findings, along with a long history of safety for protein vaccines, suggest that the REVC-128 is a safe, stable and efficacious single-shot vaccine candidate to induce the earliest protection against SARS-CoV-2 infection.

---

SARS-CoV-2, the virus causing the COVID-19 pandemic, is a new emerging virus. SARS-CoV-2 belongs to the coronavirus family with members including severe acute respiratory syndrome coronavirus (SARS, 2003 strain), Middle East respiratory syndrome (MERS) and others causing the common cold. The development of vaccine candidates focuses on the spike (S) protein of the SARS-CoV-2 virus, which forms homotrimers protruding from the virus surface and mediates virus entry by targeting angiotensin receptor 2 (ACE2) as the receptor^1^ and heparin as co-receptor. S protein is comprised of two functional subunits: S1 for receptor binding and S2 for mediating fusion of the viral and cellular membranes (Fig. 1A). For SARS-CoV-2, S protein is cleaved at the boundary (S1/S2) between S1 and S2, which remains non-covalently bound in the prefusion conformation^2^ (Fig. 1B). The S1 subunit comprises the N-terminal domain (NTD) and receptor binding domain (RBD), while S2 subunit contains the fusion machinery with fusion peptide (FP) located downstream of the cleavage site (Fig. 1A). The second cleavage at the S2’ site within the S2 subunit leads to a conformational change to initiate the membrane fusion^3^ (Fig. 1A). The discovery of neutralizing monoclonal antibodies (nAbs) reveals a number of vulnerable sites of the virus. Currently, most of the discovered nAbs have been reported to target the RBD^4–8^ and NTD^4,9^, in contrast to a small number of nAbs targeting the S2 subunit^9^. In particular, the footprint of the most potent nAbs usually lines within the epitope for ACE2 binding on the RBD, suggesting that RBD is a desirable neutralizing epitope on virus spike protein.

Vaccines with a multivalent display of antigen are believed to induce longer-lasting immunity than monovalent antigens^10,11^. Multivalent display using virus-like particle (VLP) or nanoparticle (NP) is the common strategy for vaccine development, such as the VLP comprising an array of 360 copies of the L1 capsid protein for the licensed HPV vaccine^12^, or the eOD-GT8 60mer HIV-1 vaccine currently in clinical trials^13,14^. Spike protein or RBD of SARS-CoV-2 conjugated on NP were shown to elicit potent neutralizing antibody responses^15,16^. The Novavax COVID-19 nanoparticle vaccine, NVX-CoV2373, induced protection for mice^17^ and macaques^18^ against viral challenge and showed 89.3% efficacy in a Phase 3 clinical trial conducted in the UK, by using a two-dose regimen. *Helicobacter pylori* ferritin has been used to display antigens from influenza^19,20^, hepatitis C virus^21^, HIV-1^22,23^, Epstein-Barr virus^24^ and SARS-CoV-2^25^. Ferritin is a highly conserved protein with a 24-subunit protein shell. Very recently, Powell et al.^25^ reported that ferritin display of SARS-CoV-2 spike ectodomain is able to induce a potent neutralizing antibody response in mice. Similarly, influenza ferritin vaccines have been shown to be safe in clinical trials (NCT03186781 and NCT03814720).

In this study, we developed a COVID-19 nanoparticle vaccine, designated as REVC-128 (or spike NP) with trimeric spike ectodomain subunits (glycine substitution at residue 614) multimerized onto the ferritin nanoparticle. The design of this vaccine is aimed to preferentially present the neutralizing antibody epitope (RBD) but occlude the S2 subunit to the immune system. Such design elicits the neutralizing antibody response over cross-reactive antibody, which might minimize antibody-dependent enhancement (ADE) concern (see Discussion section). We compared the immunogenicity of spike NP versus spike non-NP (soluble trimeric spike protein without nanoparticle presentation) and observed that a single dose of spike NP induced significantly higher neutralizing but less S2 subunit-specific or cross-reactive antibody titers than spike non-NP in mice. Encouraged by the observation of a high neutralizing antibody titer (10^4^ ID_50_ of serum dilution) induced by spike NP at two weeks post immunization, we sought to evaluate the protection efficacy of one-dose regimen with virus challenge. The in-vivo protection efficacy study in hamsters showed that vaccinated animals slightly gained body weight from 4 days post infection, while the sham group lost ~10% weight by 7 days post infection. To our knowledge, REVC-128 is the first COVID-19 vaccine to show evidence of vaccine induced-protection starting at two weeks post immunization in this virus challenge model, which is earlier than other vaccine candidates showing induced-protection starting at or after four weeks post first dose of immunization (see Discussion section).

**Figure 1.**
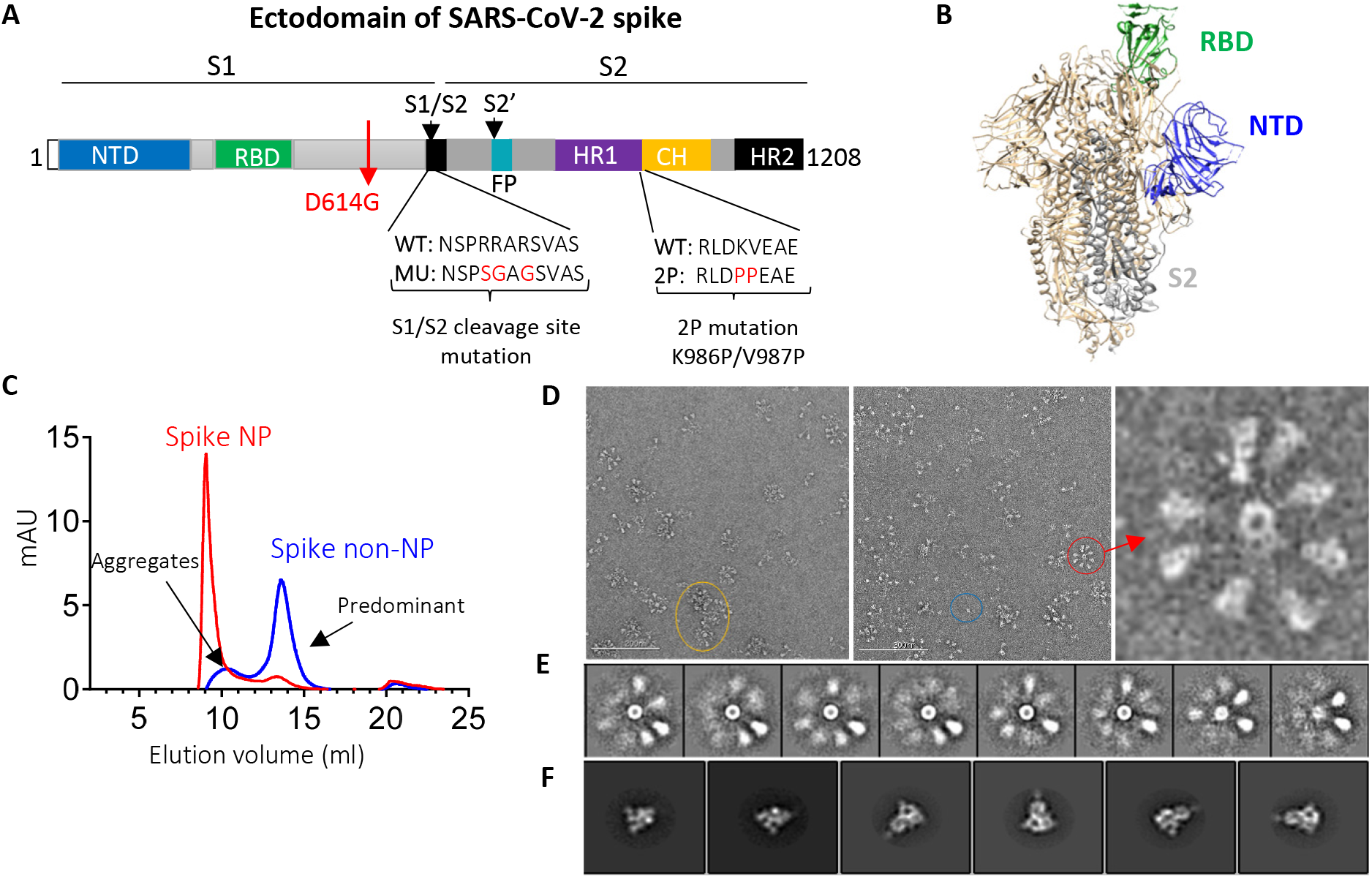
SARS-CoV-2 spike ectodomain and nanoparticle presenting trimeric spike ectodomain. (**A**) Schematic of SARS-CoV-2 spike protein ectodomain. NTD: N-terminal domain; RBD: receptor-binding domain; S1/S2 = S1/S2 protease cleavage site; FP = fusion peptide; HR = heptad repeat. Two arrows indicate the cleavage sites. The native furin cleavage site was knocked out (RRAR→SGAG), two proline at positions K986 and V987 substituted, and one glycine at position D614 substituted for ectodomain expression and nanoparticle conjugation. (**B**) Schematic of prefusion conformation of SARS-Cov-2 trimeric S structure with NTD, RBD and S2 subunit highlighted in blue, green and grey on one protomer, respectively (PDB:6VSB). (**C**) Size-exclusion chromatography (SEC) profiles of spike NP (red) and spike non-NP (blue) presentation on a Superose 6 column. (**D**) Spike NP observation by negative stain EM. In the raw micrograph, the representative of nanoparticle single particle, spike NP aggregates and NPs with varying stoichiometries was circled in blue, yellow and red, respectively. The closer observation of a selected multivalent spike NP is on the right. The grey scale bar represents 200 nm. (**E**) 2D classes averages of spike NP. The pictures show varying numbers of spike proteins on NPs. (**F**) Spike trimers are in the desired prefusion conformation on NP.

## Results

### Generation of trimeric spike protein with or without nanoparticle presentation

We first expressed SARS-CoV-2 spike ectodomain residues 1 to 1208 in trimeric form by appending a T4 fibritin trimerization motif to the c-terminus of spike ectodomain. The ectodomain contains a glycine substitution at residue 614 to match viral predominant isolate circulating in the middle of 2020^26–28^, a “SGAG” substitution at the furin cleavage site (residues 682-685) to knockout furin cleavage, and two proline substitutions at residues 986 and 987 to increase stability^29^ (Fig. 1A). The trimeric ectodomain protein was further multimerized onto ferritin with a linker to generate a nanoparticle (NP) presenting trimeric spike protein. Trimeric spike proteins with or without NP presentation were referred to spike NP (also designated as REVC-128) or spike non-NP, respectively in the following. We first characterized spike NP or spike non-NP on size-exclusion chromatography (SEC) with overlapping profiles showing that spike NP (red) was significantly larger than spike non-NP (blue) (Fig. 1C). Spike NP displayed a clear sharp peak, while spike non-NP displayed two peaks that we assigned to a minor aggregates peak and a predominant trimer fraction peak (Fig. 1C). Negative stain electron microscopy (nsEM) was used to further evaluate the conformational integrity of spike NP proteins. Imaging of spike NP revealed the forms of single particles (blue circled), spike NP aggregates (yellow circled) and spike NPs with varying stoichiometries (red circled), with the latter being the most predominant (Fig. 1D). The majority of stoichiometries ranged from 2-9 spike proteins with one representative particle shown in Fig. 1D. Closer evaluation of spike proteins further validated the order and pre-fusion homogeneity of the spikes on the NPs (Figs. 1E and F). Consistent with our vaccine design, these nsEM observations validated that the arrangement of spike proteins on the NP sterically blocks S2 subunits by the proximity of adjacent spikes (Fig. 1B), and this blockage depends on the occupancy rate of the spikes on the NP.

### In-vitro characterization and comparison of spike NP and non-NP

Ideally, trimer mimetics of the native spike on NP or by itself should present all epitopes recognized by the neutralizing antibodies (nAbs). To characterize the antigenic profile of spike trimers, both spike NP and spike non-NP were tested to bind to a panel of published nAbs (IgG format) targeting the RBD and NTD^4,9,30,31^, a non-neutralizing antibody CR3022^32^ and an HIV antibody as negative control in ELISAs. The binding of spike NP or non-NP to all tested IgGs were potent, with the exception of the HIV antibody control (Fig. 2A). We next sought to compare the binding kinetics of two representative nAbs to spike NP versus non-NP by Bio-Layer Interferometry (BLI). To eliminate the multivalent binding on BLI, we first generated antibody Fab using sequences from nAbs, COVA1-18 (RBD-specific) and COVA1-22 (NTD-specific)^4^. Fabs of COVA1-18 and COVA1-22 were immobilized on anti-human Fab-CH1 sensors and probed with spike NP or non-NP at 7 different concentrations. BLI data showed that both Fabs binding to spike NP had higher affinities (<pM level), compared to spike non-NP (3.54 and 0.033 nM for COVA1-18 and COV1-22 Fabs, respectively), and this higher affinity to spike NP was attributed to the slower dissociation off-rates (Fig. 2B). Compared to NTD-specific antibody COVA1-22 binding, the difference of binding to RBD-specific antibody COVA1-18 between spike NP and non-NP was more striking (Fig. 2B), suggesting that spike NP exhibits the RBD more robustly. The overall antigenic profile determined by ELISA and BLI (Fig. 2) confirmed that spike NP and non-NP all displayed the favorable epitopes targeted by the tested nAbs.

**Figure 2.**
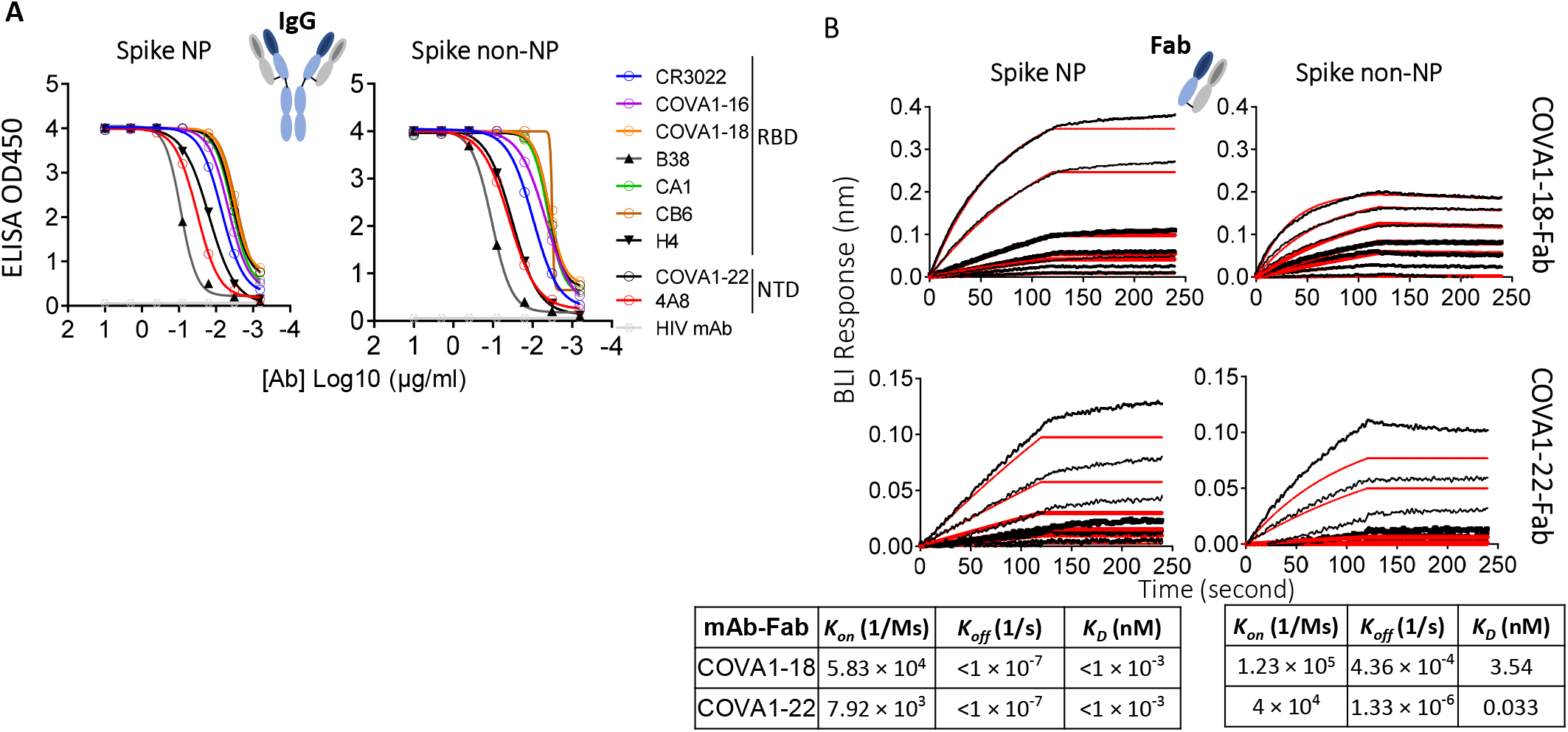
Characterize and compare antigenicity of spike NP and spike non-NP. (**A**) Antibody (IgG format) binding to spike NP protein (left) and spike non-NP (right) in ELISAs with raw curves displayed. Antibodies targeting RBD and NTD were indicated with a control HIV antibody. (**B**) Kinetics of antibody Fab-spike binding characterization by Bio-Layer Interferometry (BLI). BLI curves were generated with two published antibody Fab format COVA1-18 on the top and COVA1-22 at the bottom, immobilized on anti-human Fab-CH1 sensors, followed by probing with spike NP or non-NP proteins at concentrations of 250, 125, 62.5, 31.3, 15.6, 7.8 and 3.9 nM. Raw and fit curves were labeled in black and red, respectively. Binding kinetic measurements were indicated below the sensograms.

Spike NP was initially stored at −80°C. To understand the stability of spike NP stored at different temperatures, we evaluated the conformational integrity and antigenicity of this protein when stored at 4°C, 22°C (room temperature, RT), and 37°C for a period of 2 days, 1 week and 4 weeks. Three representative antibodies targeting the RBD, NTD and S2 subunits of SARS-CoV-2 spike protein, and a negative control antibody were used to test binding. As shown in Fig. S1, spike NP stored at different temperatures for up to 4 weeks displayed binding to selected RBD and NTD-specific antibodies-CB6 and 4A8^9,31^ respectively, identical to positive control spike NP protein stored at −80°C, although we observed slightly decreased binding to the S2 subunit-specific antibody, RV82. RV82 is a fully human monoclonal antibody (IgG1) that was identified from a B cell repertoire of COVID-19 convalescent human and confirmed to bind to monomeric or trimeric S2 subunit proteins of SARS-CoV-2 (Fig. S2A). Consistently, RV82 retained binding to trimeric spike protein of SARS-CoV-2 B.1.351 variant (initially identified in the South Africa) that has all mutations located within S1 subunit, while most of tested RBD- or NTD-specific antibodies showed ablated binding to these mutations, with the exception of CR3022^32^ and COVA1-16^4^ antibodies (Left two, Fig. S2B). Different from CR3022 and COVA1-16, RV82 bound to SARS-CoV-2, but not SARS (2003 strain) (Right, Fig. S2B). Together with the notion that RBD and NTD are the major neutralizing epitopes^4,5,9,31^, and SDS-PAGE analysis of spike NP showed no difference between protein being stored at various temperatures and −80°C (data not shown), we suggest that spike NP is stable up to 37°C for at least 4 weeks. In the future, other methods to verify the thermostability, such as EM and DSF (Differential Scanning Fluorimetry), will be utilized.

### Immunogenicity of spike NP and non-NP in mice

We next evaluated the immunogenicity of spike NP and non-NP in mice. Two groups of mice were immunized with spike NP and spike non-NP with the Sigma Adjuvant System via subcutaneous injection route, respectively. A third group of mice were injected with PBS as a negative control. We first assessed antibody binding of sera collected 14-or 28-days post immunization to trimeric spike protein with D614G mutation. Significant levels of spike protein-specific IgG were detected in all vaccinated mice 14 days post immunization and spike NP induced spike-specific IgG ~1.5-fold higher than spike non-NP on days 14 and 28, with the titer declining on day 28 (Fig. 3A). Besides trimeric spike protein binding, we assessed the binding of RBD, S2, and NTD subunits to sera collected 14 days post immunization. The results showed that sera from spike NP immunized mice displayed significantly higher binding to RBD than sera from spike non-NP immunized animals (** *p*<0.01, Mann-Whitney test) (Left, Fig. 3B), while serum binding to S2 subunit was opposite, with sera from spike non-NP immunized animals showing significantly stronger binding to S2 subunit (** *p*<0.01, Mann-Whitney test) (Middle, Fig. 3B), indicating NP presentation preferentially exposed RBD over S2 subunit, consistent with vaccine design. NTD-specific antibody responses in all groups were weak on day 14 (Right, Fig. 3B).

**Figure 3.**
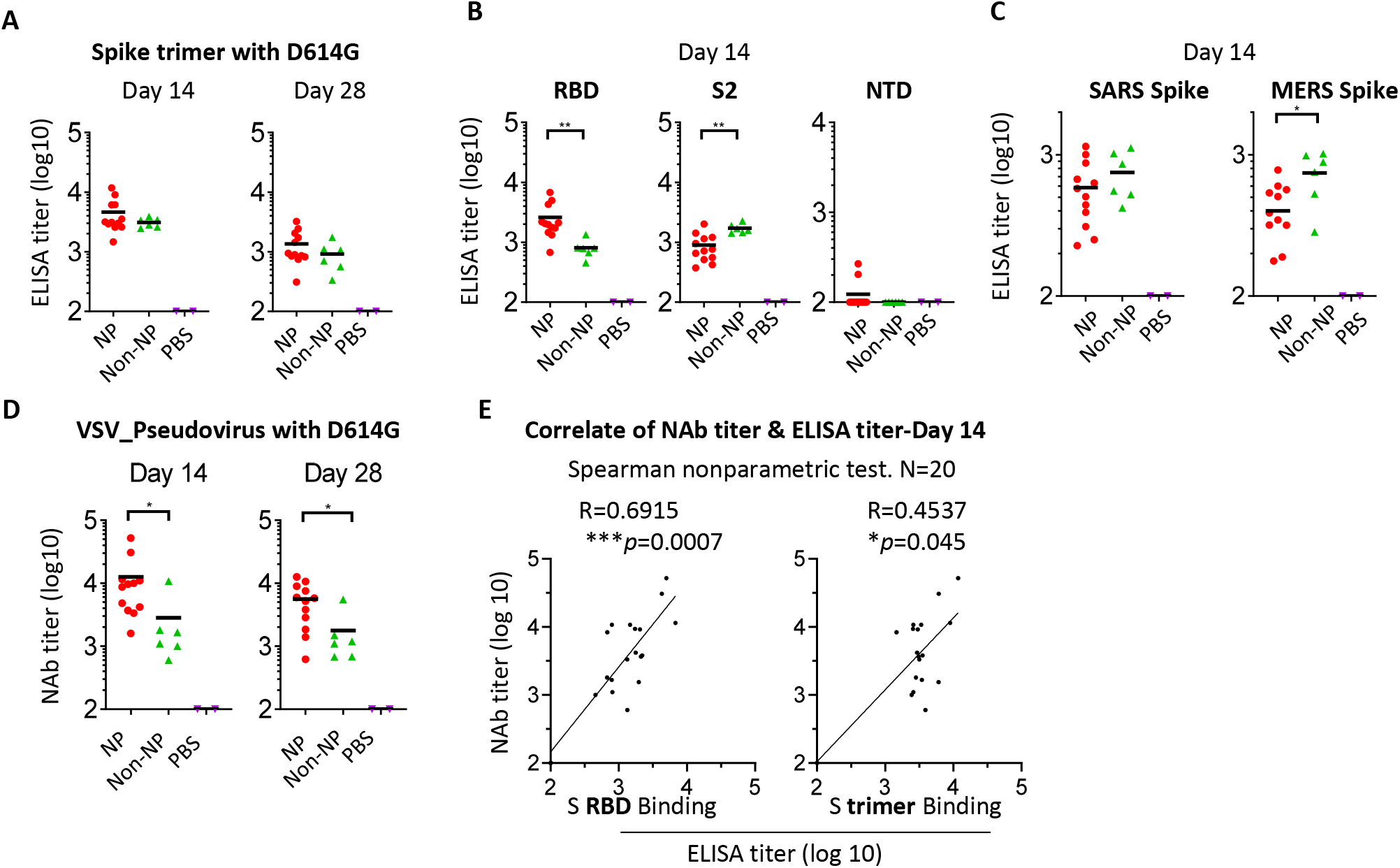
Immune response to spike NP or non-NP. (**A**) Wide-type C57BL/6 mice were immunized with 20 μg spike NP or spike non-NP with Sigma Adjuvant System via subcutaneous injection route. The serum was collected 14-and 28-days post immunization and tested to bind to SARS-CoV-2 spike trimeric protein with D614G mutation in ELISAs. ELISA titer was calculated on reciprocal serum dilution to achieve 50% of maximal optical absorbance (OD). Black bars reflect mean responses. (**B**) The binding of sera collected at day 14 to spike RBD, S2 and NTD subunits of SARS-CoV-2 in ELISAs. Statistical analysis was performed with Mann-Whitney test (** *p*<0.01). (**C**) The binding of sera collected at day 14 to trimeric spike protein of SARS (2003 strain) and MERS. Statistical analysis was performed with Mann-Whitney test (* *p*<0.05). (**D**) The neutralizing activity of sera collected at days 14 and 28 against VSV pseudotyped virus with SARS-CoV-2 spike protein containing D614G mutation. NAb titer (neutralizing antibody) represents the reciprocal of the antiserum dilution at which virus entry is inhibited by 50%, when calculated after curve-fitting with the Prism program (GraphPad). Black bars reflect mean responses. Statistical analysis was performed with Mann-Whitney test (* *p*<0.05). (**E**) The correlate of serum neutralizing titer and ELISA titer of binding to RBD protein (left) or trimeric spike (right). The correlation for day 14 sera between neutralizing titer (log10) and ELISA binding titer (log10) was analyzed using Spearman nonparametric test. Line represents the best fit linear regression.

Cross-reactive and non or weak neutralizing antibodies are potentially responsible for antibody-dependent enhancement (ADE). To evaluate vaccine-elicited cross-reactive antibodies, we assessed sera collected on day 14 for binding to trimeric spike proteins from SARS (2003 strain) or MERS. Consistent with S2 subunit binding, sera from spike non-NP immunized animals showed stronger binding to these two different coronavirus spike proteins than sera from spike NP immunized ones, especially to the MERS spike protein (* *p*<0.05, Mann-Whitney test) (Fig. 3C). The data suggested spike NP elicited less cross-reactive antibodies that are possibly S2 subunit-specific.

Neutralizing antibodies are related to vaccine-induced protection. Sera collected on days 14 and 28 were tested for their neutralizing antibody activity against VSV pseudotyped virus with SARS-CoV-2 spike protein containing D614G mutation and a luciferase reporter. On day 14, we observed that the neutralizing antibody (nAb) titer of sera from spike NP immunized mice was on average 4 log (ID_50_), significantly higher than the titer of sera from spike non-NP immunized ones (* *p*<0.05, Mann-Whitney test) (Left, Fig. 3D). Similarly, the neutralizing antibody titer declined on day 28, but the titer of spike NP immunized sera was still significantly higher than that of spike non-NP immunized sera (* *p*<0.05, Mann-Whitney test) (Right, Fig. 3D). To assess neutralizing antibody epitope on spike protein, we analyzed the correlation between day 14 serum neutralizing titer and ELISA titer obtained from either binding to spike RBD subunit or whole spike protein. This analysis revealed a more correlated relationship between neutralizing titer and RBD binding titer (R = 0.6915, *** *p* = 0.0007), compared to the correlate with spike binding (R = 0.4537, * *p* = 0.045) (Fig. 3E), indicating RBD is the major neutralizing antibody epitope and RBD-specific nAbs are responsible for serum neutralizing activity. This is in agreement with the rationale of our vaccine design by using NP to preferentially present RBD, validated by the above BLI results (Fig. 2B).

### Protection efficacy of spike NP in hamsters

Encouraged by the potent immune mouse serum neutralizing antibody response elicited by spike NP that we observed on day 14, we next evaluated vaccine protection efficacy in Syrian golden hamsters, one of few small animal models susceptible to infection by the SARS-CoV-2 virus^33,34^. Two groups of hamsters were immunized with a single dose of spike NP (REVC-128) or sham control including same NP presenting Marburg trimeric GP, and same adjuvant via intramuscular injection route (Fig. 4A). We observed that sera collected from spike NP immunized hamsters on day 13, one day prior to virus challenge, displayed significantly higher potent neutralizing antibody activity against VSV pseudotyped with SARS-CoV-2 D614G spike than sera from the sham group (* *p*<0.05, Mann-Whitney test) (Fig. 4B).

**Figure 4.**
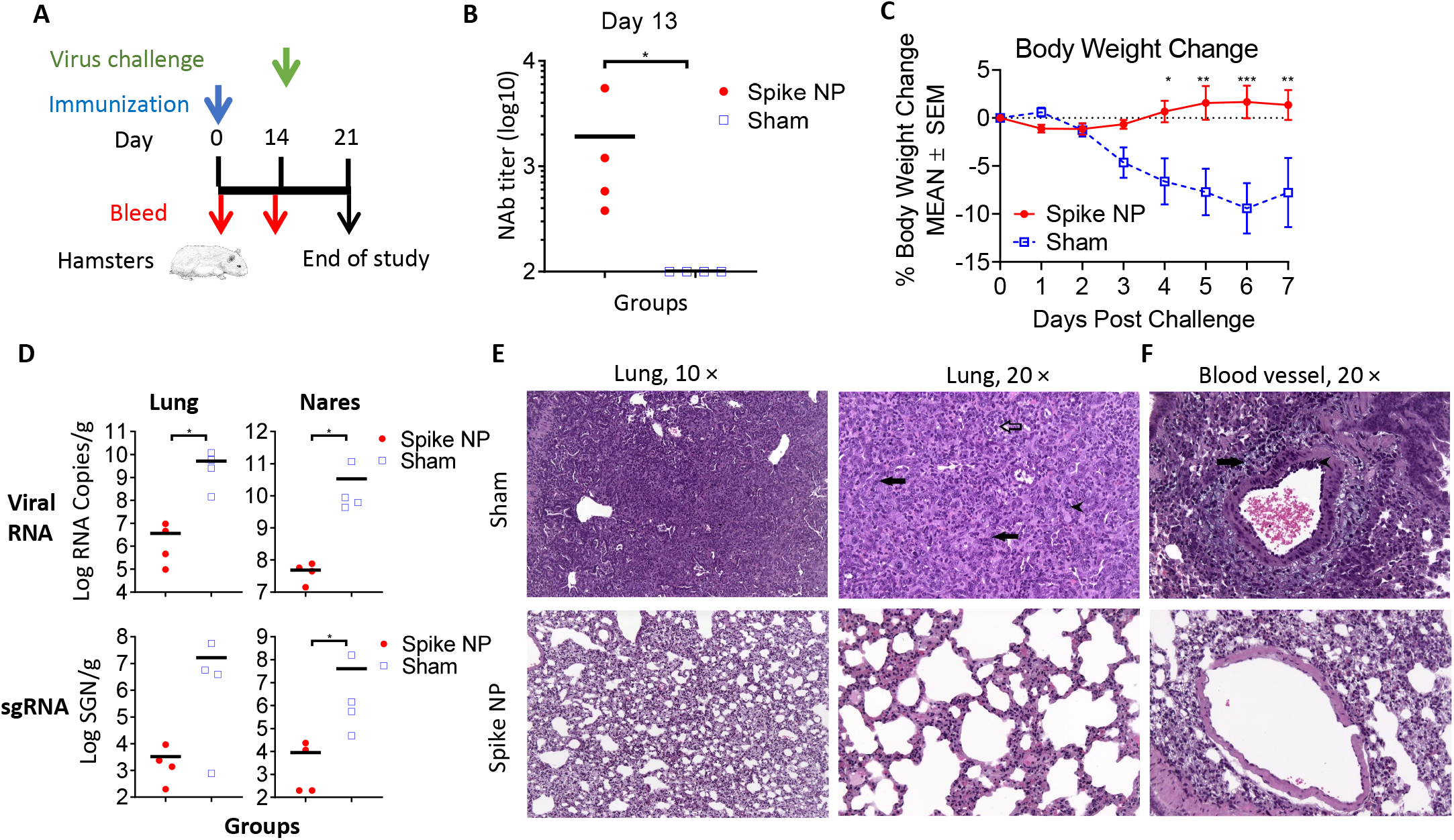
Vaccine protection efficacy against virus challenge in hamsters. (**A**) Schematic of the immunization and virus challenge protocol. Syrian golden hamsters (2F/2M) were immunized with 100 μg spike NP (REVC-128) or sham with Sigma Adjuvant System via intramuscular injection route, and challenged with 1.99 × 104 TCID50 of SARS-CoV-2 virus (USA-WA1/2020, NR-53780, BEI Resources) by the intranasal route at day 14 post immunization. (**B**) The neutralizing activity of sera collected at day 13 post immunization against VSV pseudotyped virus containing SARS-CoV-2 spike with D614G mutation. Statistical analysis was performed with Mann-Whitney test (* p<0.05). (**C**) Median percent weight change after challenge. Statistical analysis for body weight change was performed for comparison between spike NP and mock immunized animals by two-way ANOVA test, * p<0.05, ** p<0.01, *** p<0.001. (**D**) Tissue viral loads on 7 dpi. Viral loads of lung (left) and nares (right) were measured by RT-PCR and quantitated as total viral RNA copies per gram tissue (upper) and subgenomic N RNA copies per gram tissue (bottom). Limitation of quantification is 200 copies/g. Black bars reflect mean responses. Statistical analysis was performed with Mann-Whitney test (* p<0.05). SGN=Subgenomic N RNA copies. (**E, F**) Representative images of histopathology for lungs (**E**) and blood vessels (**F**) of sham control (upper) and spike NP (bottom) immunized animals. In a higher magnification of (**E**) on right (20 ×), black arrows indicate bronchiolo-alveolar hyperplasia characterized by hyperplastic epithelial cells extending from bronchioles and lining alveoli. Black arrowhead indicates hyperplastic cells with enlarged nuclei. Open arrow indicates mixed cell inflammation observed in alveolar lumen. In (**F**), black arrow indicates expansion of surrounding vascular tissue by edema (increased clear space and a pale basophilic material) and mononuclear cells. Black arrowhead indicates mononuclear inflammatory cells expanding the vessel wall (tunica media and intima).

Fourteen days post immunization, animals were challenged with SARS-CoV-2 virus intranasally. In the sham control, hamsters lost a median of 10% body weight by 7 days post infection (dpi), while spike NP immunized animals gained body weight lightly from 4 dpi (Fig. 4C). Significant weight differences post infection were observed from 4-7 dpi (* *p*<0.05, ** *p*<0.01, *** *p*<0.001, two-way ANOVA test) (Fig. 4C). To determine the impact of vaccine-induced immunity on tissue viral load, we measured both viral genomic and subgenomic RNA amounts in lung and nares collected on 7 dpi. Viral genomic RNA (vRNA) reflects remaining viral inoculum plus newly replicating virus, while subgenomic RNA (sgRNA) should only result from newly replicating virus^35^. For hamsters immunized by spike NP, we observed the significantly lower amounts of viral RNA in lung and nares (~3 log) than that in animal tissues in the sham control (Upper, Fig. 4D). Similarly, the level of sgRNA in protected animals’ tissues was approximately 3 log lower than that in sham control animals’ tissues (Bottom, Fig. 4D).

We next performed histopathology analyses of the lungs and nares of two groups of infected hamsters. Qualitative and semiquantitative analyses of lung and nares tissues collected on 7 dpi were performed using a severity scoring scale^36,37^ (Table 1). In the sham control, all hamsters showed evidence of pulmonary lesions with the observations of moderate to marked bronchiolo-alveolar hyperplasia (grade 3 or 4), mild mixed cell bronchiolo-alveolar inflammation (grade 2), and minimal to mild mononuclear cell perivascular/vascular infiltrate or alveolar/perivascular/vascular inflammation (grade 1 or 2) (Figs. 4E, F and Table 1). Perivascular edema (grade 1 or 2), syncytial cells and hemorrhage (grade 1) were observed in lungs of some but not all animals in the sham control (Fig. 4F and Table 1). The majority of animals in the sham control also exhibited atypia of hyperplastic alveolar epithelial cells with hypertrophic and variably shaped nuclei and prominent nucleoli (Fig. 4E). Syncytial cells, enlarged multinucleated cells, most likely Type II pneumocytes demonstrating viral cytopathic-like changes, were scattered throughout alveoli in lungs of animals in sham control. SARS-CoV-2 virus has been reported to infect both Type I and Type II pneumocyte cells in animal studies^38^, and the infection and loss of Type I pneumocytes would contribute to the proliferation of Type II pneumocytes. Overall, the histological observations in the lungs and nares of sham control were consistent with SARS-CoV-2 infection reported previously^39^. In spike NP immunized animals, one animal showed minimal bronchiolar hyperplasia in the lung (grade 1) and another one showed minimal lesions in nares (grade 1) (Table 1), while the rest of animals in this group showed negligible histopathologic phenotype.

**Table 1.**
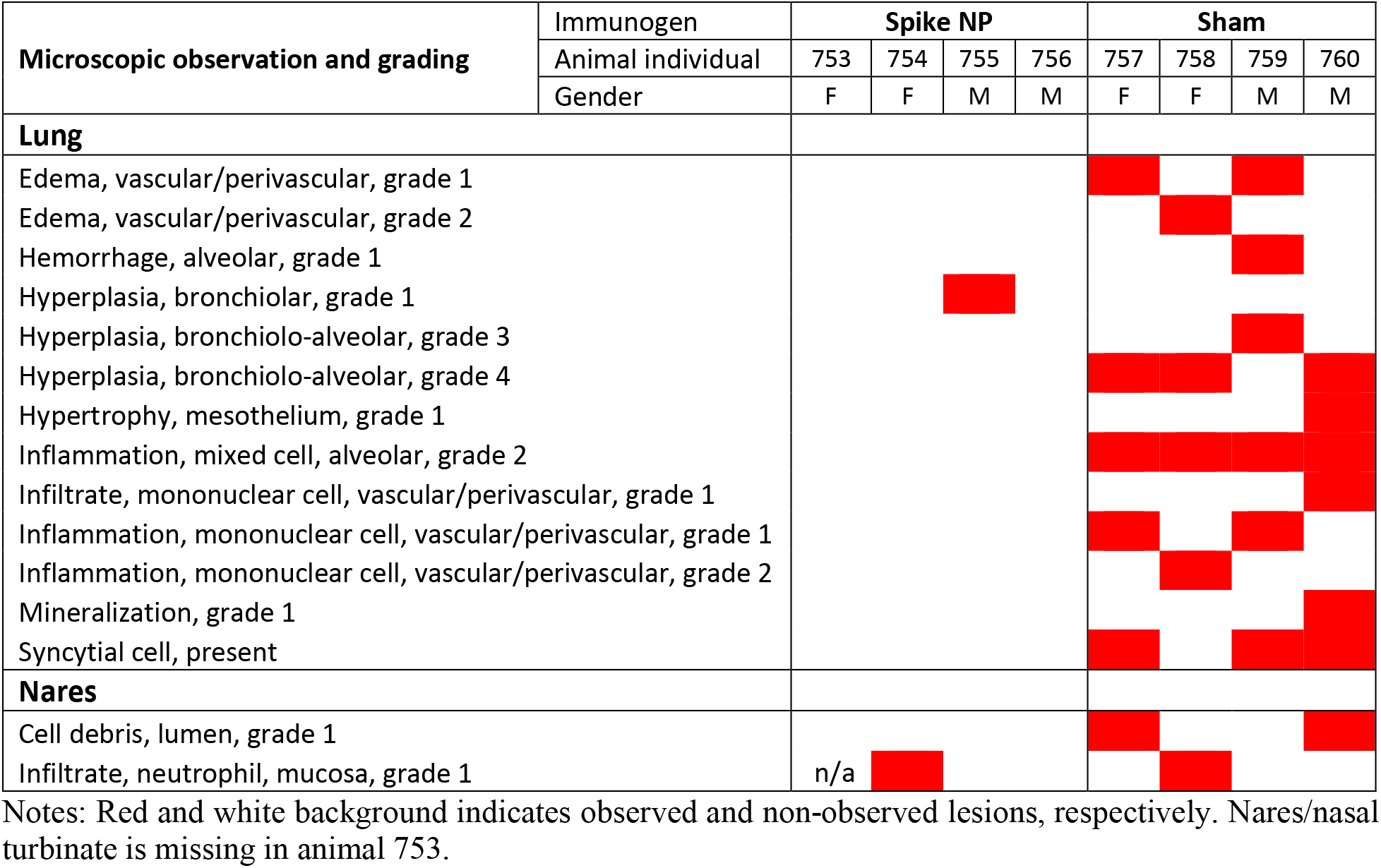
Macroscopic observation of lung and nares from spike NP immunized and sham control on 7 dpi.

In summary, weight change, viral load and histopathological data support that REVC-128 effectively protected against COVID-19 associated morbidity starting two weeks post a single dose of immunization.

## Discussion

Herein, we developed a COVID-19 protein-based nanoparticle vaccine, REVC-128. Protein-based vaccines, such as the Hepatitis B vaccine that has been administered on the first day of life for newborn babies, have shown minimal safety concern. The backbone of the nanoparticle, ferritin, has also been shown to be safe for influenza vaccines in clinical trials. Therefore, REVC-128 has the potential to protect individuals who are medically unable to receive other COVID-19 vaccines, such as people who are allergic to mRNA vaccines. Additionally, the vaccine stability data (Fig. S1) support a much less stringent requirement for the storage of this vaccine, compared with mRNA vaccines that require −20°C or −80°C storage. Vaccine storage and transportation in ambient temperature condition will enable broader availability of such vaccine.

Vaccination using a one-dose regimen enables easier deployment, tracking and administration, compared to a two-dose regimen. Further, vaccine with the capability to induce early protection can more efficiently prevent viral spread and contribute to the quelling of this COVID-19 pandemic. Herein, we showed that a single dose of REVC-128 provides protection starting two weeks post immunization, earlier than other vaccine candidates. To our knowledge, two viral vector vaccines (VSV and Ad26) have been reported to induce early protection starting four weeks post immunization using a one-dose regimen^33,40^, while mRNA vaccines (e.g., Moderna, Pfizer/BioNTech), inactivated vaccine (e.g., Sinovac Life Sciences) and another nanoparticle protein vaccine, Novavax’s NVX-CoV2373 that all use a two-dose regimen with evidence of protection occurring at or after four weeks post the first dose of immunization. Although viral vector vaccines induced early protection, the preexisting immunity to viral vectors presents a formidable challenge for this platform, especially when a boosting immunization is required. The booster enhances the immune response to viral vector, which could impair vector entry into host cells. In contrast, protein or mRNA vaccines avoid this challenge and allow more flexibility for multiple boosts to achieve a longer protection period or protection against mutant variants. We observed the decreased neutralizing antibody titer on day 28 after a single dose of immunization (Fig. 3D), and also observed the enhanced titer following a booster (data not shown). The protection durability induced by both one-dose and two-dose regimens merits further investigation. The immunogen dose used in this report is high (20 μg per mouse or 100 μg per hamster), warranting a dose-deescalating study to assess the optimal dose in the future.

During this COVID-19 pandemic, concerns arose about pre-existing human coronavirus-specific antibodies generated during previous infections. These antibodies may mediate antibody-dependent enhancement (ADE), worsening symptoms when patients are infected with SARS-CoV-2^41^. This might be one of the reasons why older adults are at higher risk for severe disease, as antibodies previously generated in response to common human coronaviruses in the elderly facilitate SARS-CoV-2 entrance into target cells, leading to more severe symptoms^42,43^. On the contrary, seroprevalence of community-acquired coronavirus in pediatrics is not common^44^. In addition, one possible explanation for SARS-CoV-2 reinfection is the antibodies induced by the first infection may help, rather than fight, the second infection, which is linked to ADE^45^. Previous Dengue virus vaccine studies revealed human clinical safety risk related to ADE^46^, resulting in vaccine trial failure. The envelope protein sequence alignment of four serotypes of Dengue and Zika viruses (flavivirus family) indicates that envelope protein domain II containing fusion machinery exhibits a higher degree of homology among these viruses, and virus infection induces high level of domain II-specific antibodies that are non or weakly neutralizing, but responsible for ADE effect^47–49^. In line with Dengue domain II, the S2 subunit of the SARS-CoV-2 spike protein containing fusion machinery exhibits a higher degree of homology among coronaviruses than the S1 subunit of spike protein. It has been reported that antibodies from SARS-CoV-2 naïve donors who had reactivity to seasonal human coronavirus strains (such as OC43 and HKU1) were cross-reactive against the nucleocapsid and S2 subunit on spike protein of SARS-CoV-2^50^. More studies have reported that cross-reactive mAbs largely target this more conserved S2 subunit on spike protein^51,52^. Conceptually, a number of S2 subunit-specific antibodies is non or weakly neutralizing, and potentially responsible for ADE. Thus, vaccine design to minimize the elicitation of S2 subunit-specific antibodies should be considered, especially when/if ADE will be observed for SARS-CoV-2.

Although ADE has yet to be fully observed for SARS-CoV-2 infection or vaccination, previous coronavirus vaccine candidates were reported to be complicated by ADE. A viral vector vaccine of the original SARS was found to enhance the immunopathology of immunized animals following viral challenge, resulting in a strong inflammatory response and even lung injury^53,54^. Mice immunized by Ad5 viral vector expressing MERS vaccine were also reported to exhibit pulmonary pathology following viral challenge, despite the vaccine conferring protection^55^. Similar observation was reported for inactivated virus vaccine. Lung immunopathology was observed when animals were immunized with inactivated whole-virus MERS vaccine, followed by virus challenge^56^. One strategy to offset ADE concern of Dengue and Zika vaccines is to reduce cross-reactive antibody response^57^. Similarly, we used adjacent spike proteins on NP to sterically block the S2 subunit exposure, which was validated with evidence of less S2 subunit and cross-reactive antibodies elicited by spike NP, compared with vaccine without NP (Figs. 3B and C). The epitope mapping of these cross-reactive antibodies will be performed in the future to assess whether they are S2-specific. Nevertheless, such design might prevent the development of severe symptoms if patients are infected with other coronaviruses post-immunization, such as seasonal coronaviruses (e.g., common cold virus), or mutated SARS-CoV-2 variants.

## Materials and Methods

### Protein expression and purification

The ectodomain (residues 1-1208) of spike protein of SARS-CoV-2 was modified based on GenBank sequence of MN908947, including a glycine substitution at residue 614, a “SGAG” substitution at the furin cleavage site (residues 682-685) and two proline substitutions at residues 986 and 987. A C-terminal T4 fibritin trimerization motif, an HRV3C protease cleavage site, an 8 × His Tag and a TwinStrep Tag were conjugated with ectodomain of spike protein. Ectodomain of spike protein was also conjugated with ferritin nanoparticle (NP) with a linker to generate spike NP. The sequence was cloned into the mammalian expression vector pCAGGS. The trimeric ectodomain of spike protein of SARS-CoV-2 South African B.1.351 variant was constructed in the same way, with the exception of the following mutations^58^: L18F, D80A, D215G, L242-244del, R246I, K417N, E484K, N501Y, D614G, and A701V. The trimeric ectodomain of spike protein of SARS (2003 strain) was modified based on GenBank sequence of AY278554, including two proline substitutions at residues 968 and 969^59^, same trimerization motif, HRV3C cleavage site and tags. The trimeric ectodomain of spike protein of MERS was modified based on GenBank sequence of JX869059, including furin cleavage site knockout, two proline substitutions at residues 1060 and 1061^60^, same trimerization motif, HRV3C cleavage site and tags.

To express trimeric S2 subunit of spike protein, residues 686-1208 of SARS-CoV-2 were cloned upstream of a C-terminal T4 fibritin trimerization motif, an HRV3C protease cleavage site, an 8 × His Tag and a TwinStrep Tag. Residues 319-541 of SARS-CoV-2 were cloned with C-terminal 6 × His Tag for RBD. Similarly, residues 14-305 of SARS-CoV-2 were cloned with C-terminal 6 × His Tag for NTD.

These expression vectors were codon optimized and confirmed by sequencing prior to being transiently transfected into FreeStyle™ 293F cells (Thermo Fisher). Protein was purified from filtered cell supernatants using StrepTactin resin (IBA) or cOmplete His-Tag Purification Resin (Roche) or Jacalin (Thermo Fisher). The purified protein was subjected to additional purification or analysis by size-exclusion chromatography using a Superose 6 column.

Plasmids encoding the heavy and light chains of CR3022, COVA1-16, COVA1-18, COVA1-22, B38, CA1, CB6, H4, 4A8 and RV82 in a human IgG1 expression vector^61,62^ were transiently transfected into FreeStyle™ 293F cells and purified as described previously^62,63^. To express antibody Fab, the heavy chain variable domain was inserted into Fab expression vector containing a 6 × His Tag as previously described^63^, followed by co-transfection with light chain expression vector. Fab was purified from cell culture supernatant by cOmplete His-Tag Purification Resin (Roche).

### ELISA binding assays

Proteins of trimeric spikes of SARS, MERS, or SARS-CoV-2 or RBD, NTD and S2 subunits of SARS-CoV-2 were coated onto 96-well Maxisorb ELISA plates at 200 ng/well diluted in PBS overnight at 4°C. On the following day, the plates were washed four times with 300 μL of 1 × PBST (0.05% Tween-20) and blocked with blocking buffer (2% dry milk / 5% fetal bovine serum in PBS) for 1 hour at 37°C. After blocking, plates were washed as described above prior to adding mAbs diluted into same blocking buffer starting from 10 μg/ml or heat-inactivated animal serum starting from 100-fold dilution with 5-fold serial dilutions for 1 hour at 37°C. After incubation, plates were washed and a 1: 5,000 dilution of Goat anti-human or anti-mouse IgG-HRP conjugate (Jackson ImmunoResearch) in PBST was added for 1 hour at room temperature. The bound mAb was detected with adding 100 μl/well of 3,3′,5,5′-Tetramethylbenzidine (TMB) substrate (Life Technologies) and incubation at room temperature for 5 min prior to the addition of 100 μl of 3% H_2_SO_4_ to stop the reaction. The optical density (OD) was measured at 450 nm.

### BioLayer interferometry

Biolayer light interferometry (BLI) was performed using an Octet RED96 instrument (ForteBio, Pall Life Sciences) as described previously^62–64^. Antibody Fab was captured onto anti-human Fab-CH1 biosensors at concentration of 10 μg/ml as ligand and the tested samples of spike NP or non-NP was diluted in 7 × 2-fold series starting from 250 nM to 3.9 nM in solution, respectively. Briefly, biosensors, pre-hydrated in binding buffer (1× PBS, 0.01% BSA and 0.2% Tween-20) for 10 min were first immersed in binding buffer for 60 s to establish a baseline followed by submerging in a solution containing ligand for 60 s to capture ligand. The biosensors were then submerged in binding buffer for a wash for 60 s. The biosensors were then immersed in a solution containing various concentrations of tested samples as analyte for 120 s to detect analyte/ligand association, followed by 120 s in binding buffer to assess analyte/ligand dissociation. Binding affinity constants (dissociation constant, K_D_; on-rate, k_on_; off-rate, k_off_) were determined using Octet Analysis software.

### VSV-spike pseudovirus production and neutralization assay

To generate SARS-CoV-2 spike VSV pseudovirus, a plasmid encoding SARS-CoV-2 spike harboring a C-terminal 18-residue truncation was transfected into pre-seeded 293T cells. Next day, transfected cells were infected with VSV(G*ΔG-luciferase) (Kerafast) at an MOI of 3 infectious units/cell. The cell supernatant containing SARS-CoV-2 pseudotyped VSV was collected at day 2 post-transfection, centrifuged to remove cellular debris, aliquoted and frozen at −80°C.

Neutralization assays using above SARS-CoV-2 pseudotyped VSV were performed as previously described^65^ with modification. This pseudovirus was first titrated with duplicate on Vero E6 cells cultured in EMEM supplemented with 10% fetal bovine serum and 100 I.U./mL penicillin and 100 μg/mL streptomycin at 37°C. The dilution of pseudovirus to achieve 1,000-fold luciferase signal higher than background was selected for neutralization assay. In neutralization assay, the heat-inactivated serum starting from 100-fold dilution with serial dilutions was incubated with diluted pseudotyped virus in EMEM for 1 hour at 37°C before infecting Vero E6 cells at 37°C, 5% CO_2_ for 1 hour. The next day, cells were lysed with Passive Lysis Buffer (Promega) for 40 minutes at room temperature with shaking before the addition of the Luciferase Activating Reagent (Promega). The luminesce was read immediately on a Molecular Devices reader. Percent neutralization was calculated based on wells containing virus only and cells only as background. Data was fit to a 4PL curve in GraphPad Prism 7.

### Negative stain Electron microscopy

Negative stain electron microscopy (nsEM) was performed as previously described^65,66^. Briefly, spike NP was added to 400 square copper mesh grids coated with carbon and stained with 2% uranyl formate. The grids were imaged on a 120keV Tecnai Spirit electron microscope using an Eagle 4k × 4k CCD camera. NP particles were manually selected from the raw micrograph stacked with a box size of 200 pixels and aligned using iterative MRA/MSA^67^. Single particles were picked with DogPicker and processed in RELION 3.0.

### Animal experiments

Animal experiments were carried out in compliance with all pertinent US National Institutes of Health regulations and approval from the Animal Care and Use Committee (ACUC) of Noble Life Sciences and Bioqual. For the immunogenicity study, 6-to 8-week-old female C57BL/6 mice (Jackson Laboratory) were inoculated subcutaneously in two sites. Each animal received a single dose of 20 μg protein immunogen in 100 μl of PBS containing 50 μl of Sigma Adjuvant System (Sigma) with the procedure of immunogen and adjuvant mixture following manufacture’s manual. For serum preparation, blood samples were collected retro-orbitally on days 0, 14 and 28. For the protection efficacy study conducted at Bioqual, 7-week-old male and female Syrian golden hamsters were inoculated intramuscularly into each hind leg. Each animal received a single dose of 100 μg protein immunogen in 200 μl of PBS containing 100 μl of the same adjuvant. For serum preparation, blood samples were collected retro-orbitally on days 0 and 13. On day 14, all animals were challenged with 1.99 × 10^4^ TCID_50_ of SARS-CoV-2 virus (USA-WA1/2020, NR-53780 BEI Resources). Virus was administered as 100 μl by the intranasal route (50 μl into each nostril). Body weights were assessed daily. All animals were sacrificed on 7 dpi for tissue analyses. Challenge studies were conducted under maximum containment in an animal biosafety level 3 facility under ACUC-approved protocol in compliance with the Animal Welfare Act and other federal statutes and regulations relating to animals and experiments involving animals.

## Quantitative RT-PCR assay for SARS-CoV-2 RNA

The amounts of RNA copies per gram tissue were measured using a qRT-PCR assay as described previously^34^. Briefly, viral RNA was extracted from lung and nares collected on 7 dpi with RNA-STAT 60 (Tel-test”B”)/chloroform, precipitated and resuspended in AVE Buffer (Qiagen). To generate a control for the amplification reaction, RNA was isolated from the applicable virus stock using the same procedure. RT-PCR assays were performed using TaqMan RT-PCR kit (Bioline, BIO-78005) with primers and probe sequences described previously^34^. The signal was compared to the known standard curve and calculated to give copies per gram (g). All samples were tested in triplicate.

## Quantitative RT-PCR assay for SARS-CoV-2 subgenomic RNA

SARS-CoV-2 subgenomic mRNA (sgRNA) was determined as described previously^34^ with modification. Briefly, above extracted RNA was first reverse-transcribed using Superscript III VILO (Invitrogen) according to the manufacturer’s instructions. A Taqman custom gene expression assay (ThermoFisher Scientific) was designed using the sequences targeting the N gene sgRNA. Reactions were performed on a QuantStudio 6 and 7 Flex Real-Time PCR System (Applied Biosystems) with following primers and probe sequences. Standard curves generated using SARS-CoV-2 N gene sgRNA pre-cloned into an expression plasmid were used to calculate sgRNA in copies per gram. All samples were tested in triplicate.

Subgenomic RNA primers:

SG-N-F: CGATCTCTTGTAGATCTGTTCTC
SG-N-R: GGTGAACCAAGACGCAGTAT
Probe: FAM/TAACCAGAA/ZEN/TGGAGAACGCAGTGGG/IABkFQ

### Histopathology

Hamsters were euthanized for necropsy on 7 dpi. Lung and nares were collected in 10% neutral buffered formalin (NBF), followed by being fixed, processed to hematoxylin and eosin (H&E) stained slides and examined by a board-certified pathologist. Qualitative and semiquantitative assessment were performed as described previously^36,37^, industry best practices^68^ and terminology for data capture were consistent with International Harmonization of Nomenclature and Diagnostic Criteria (INHAND)^69,70^.

### Statistical analysis

ELISA, nAb titer or viral load statistical analyses of comparison between spike NP and non-NP or sham immunized animal sera were performed by using the Mann-Whitney test with * *p*<0.05, ** *p*<0.01. Correlation statistical analyses between ELISA and nAb titers were performed by using the Spearman nonparametric test with * *p*<0.05, *** *p*<0.001. The statistical analysis of comparison of body weight change at each time point between animals with spike NP and mock immunized was performed by using the two-way ANOVA test with * *p*<0.05, ** *p*<0.01, *** *p*<0.001 using GraphPad Prism version 8.

## Acknowledgments

We thank Dr. Michael Diamond, Washington University, for his valuable advice. This study is partially supported by fund from The University of Maryland Strategic Partnership (MPower) and Institute for Bioscience and Biotechnology Research intramural startup fund (YL). This study is also partially supported by fund from TEDCO (the Maryland Technology Development Corporation) (YW). With the mission to facilitate technology company growth in Maryland, TEDCO enhances economic development growth through the fostering of an inclusive entrepreneurial innovation ecosystem.

## Author contributions

Investigation: M.G., C-I.C., Y.L. and Y.W.; Negative stain EM experiment: J.L.T., A.M.J. and A.B.W.; Hamster study with virus challenge and viral load measurement: J.G., M.P. and S.K.; Histopathology examination: S.W. Y.W. wrote the manuscript with input from all authors.

## Severity Grading Scale

The severity of the non-neoplastic tissue lesions is graded as follows:

### Grade 1 (1+): <u>Minimal</u>

This corresponds to a histopathologic change ranging from inconspicuous to barely noticeable but so minor, small, or infrequent as to warrant no more than the least assignable grade. For multifocal or diffusely-distributed lesions, this grade was used for processes where less than approximately10% of the tissue in an average high-power field was involved. For focal or diffuse hyperplastic/hypoplastic/ atrophic lesions, this grade was used when the affected structure or tissue had undergone a less than approximately 10% increase or decrease in volume.

### Grade 2 (2+) <u>Mild</u>

This corresponds to a histopathologic change that is a noticeable but not a prominent feature of the tissue. For multifocal or diffusely-distributed lesions, this grade was used for processes where between approximately 10% and 25% of the tissue in an average high-power field was involved. For focal or diffuse hyperplastic/hypoplastic/atrophic lesions, this grade was used when the affected structure or tissue had undergone between an approximately 10% to 25% increase or decrease in volume.

### Grade 3 (3+): <u>Moderate</u>

This corresponds to a histopathologic change that is a prominent but not a dominant feature of the tissue. For multifocal or diffusely-distributed lesions, this grade was used for processes where between approximately 25% and 50% of the tissue in an average high-power field was involved. For focal or diffuse hyperplastic/hypoplastic/atrophic lesions, this grade was used when the affected structure or tissue had undergone between an approximately 25% to 50% increase or decrease in volume.

### Grade 4 (4+): <u>Marked</u>

This corresponds to a histopathologic change that is a dominant but not an overwhelming feature of the tissue. For multifocal or diffusely-distributed lesions, this grade was used for processes where between approximately 50% and 95% of the tissue in an average high-power field was involved. For focal or diffuse hyperplastic/hypoplastic/atrophic lesions, this grade was used when the affected structure or tissue had undergone between an approximately 50% to 95% increase or decrease in volume.

### Grade 5 (5+): <u>Severe</u>

This corresponds to a histopathologic change that is an overwhelming feature of the tissue. For multifocal or diffusely-distributed lesions, this grade was used for processes where greater than approximately 95% of the tissue in an average high-power field was involved. For focal or diffuse hyperplastic/hypoplastic/atrophic lesions, this grade was used when the affected structure or tissue had undergone a greater than approximately 95% increase of decrease in volume.

