## Supplemental figure and legend for "One dose of COVID-19 nanoparticle vaccine REVC-128 provides protection against SARS-CoV-2 challenge at two weeks post immunization": Supplemental figure legend.docx

**Supplementary Figures**

**Fig. S1**. Spike NP thermostability evaluation. Antibodies targeting RBD, NTD and S2 subunits of SARS-CoV-2 spike protein, and a Zika negative control antibody were used to test binding to spike NP protein stored at 4°C, 22°C (room temperature, RT), and 37°C for a period of 2 days, 1 week and 4 weeks in ELISAs with raw curves displayed.

**Fig. S2**. RV82 antibody targets S2 subunit of SARS-CoV-2 and discriminates SARS-CoV-2 from SARS. (A) RV82 antibody binds both monomeric and trimeric S2 subunit of SARS-CoV-2 spike protein. Monomeric S2 protein was purchased from Sino Biological with catalog number of 40590-V08B. Trimeric S2 protein was expressed using a plasmid containing the sequence encoding S2 subunit and a C-terminal T4 fibritin trimerization motif (see Methods for details). (B) RV82 antibody retains binding to trimeric spike protein from South African B.1.351 variant and discriminates SARS-CoV-2 from SARS (2003 strain). WT sequence of SARS-CoV-2 spike is from Wuhan strain. Antibodies targeting RBD, NTD and S2 were indicated. ACE2-Fc chimera was purchased from GenScript with catalog number of Z03484.
