## Supplementary figures and images for "One dose of COVID-19 nanoparticle vaccine REVC-128 provides protection against SARS-CoV-2 challenge at two weeks post immunization"

### Fig. S1.TIF

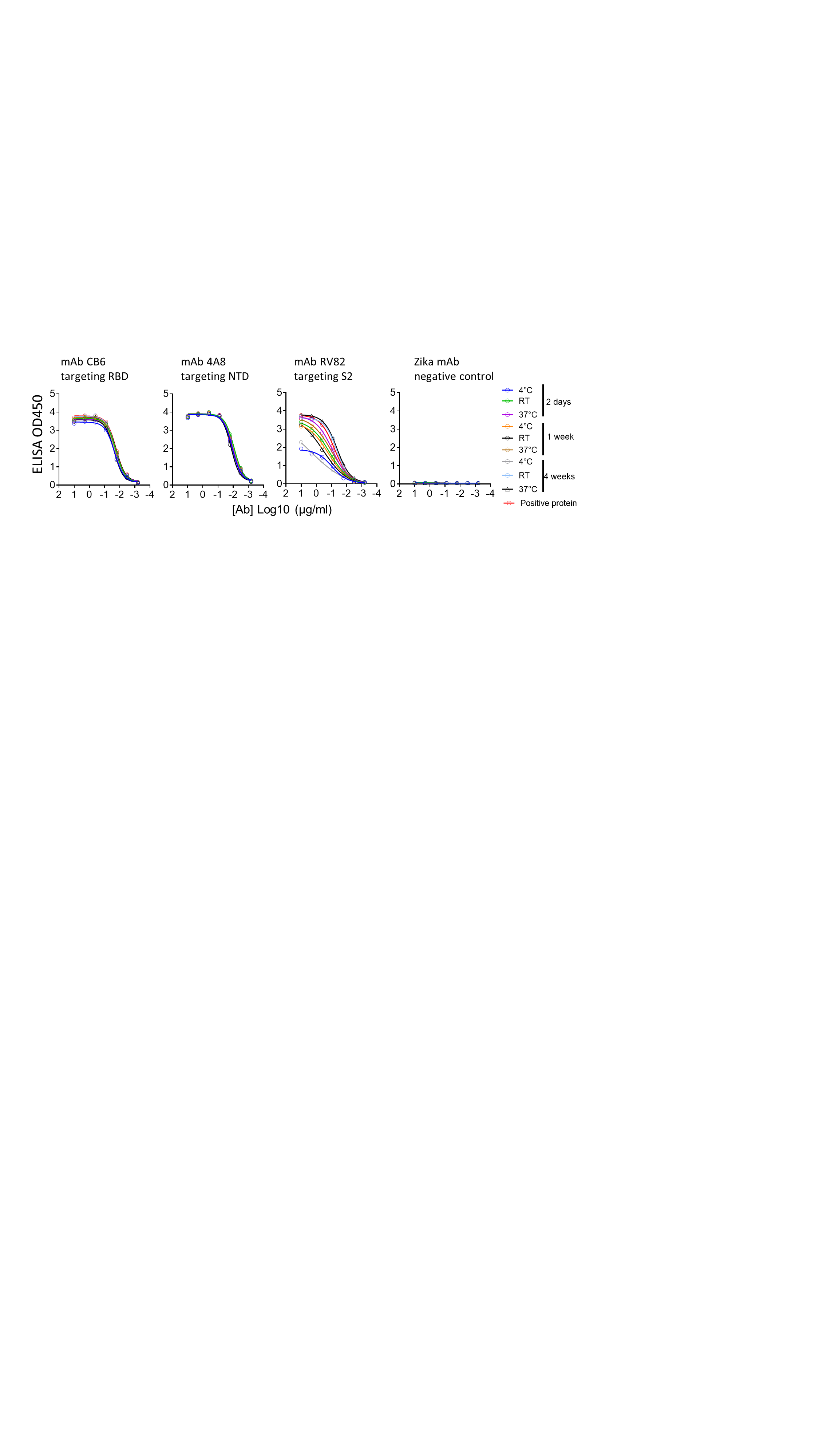

### Fig. S2.TIF

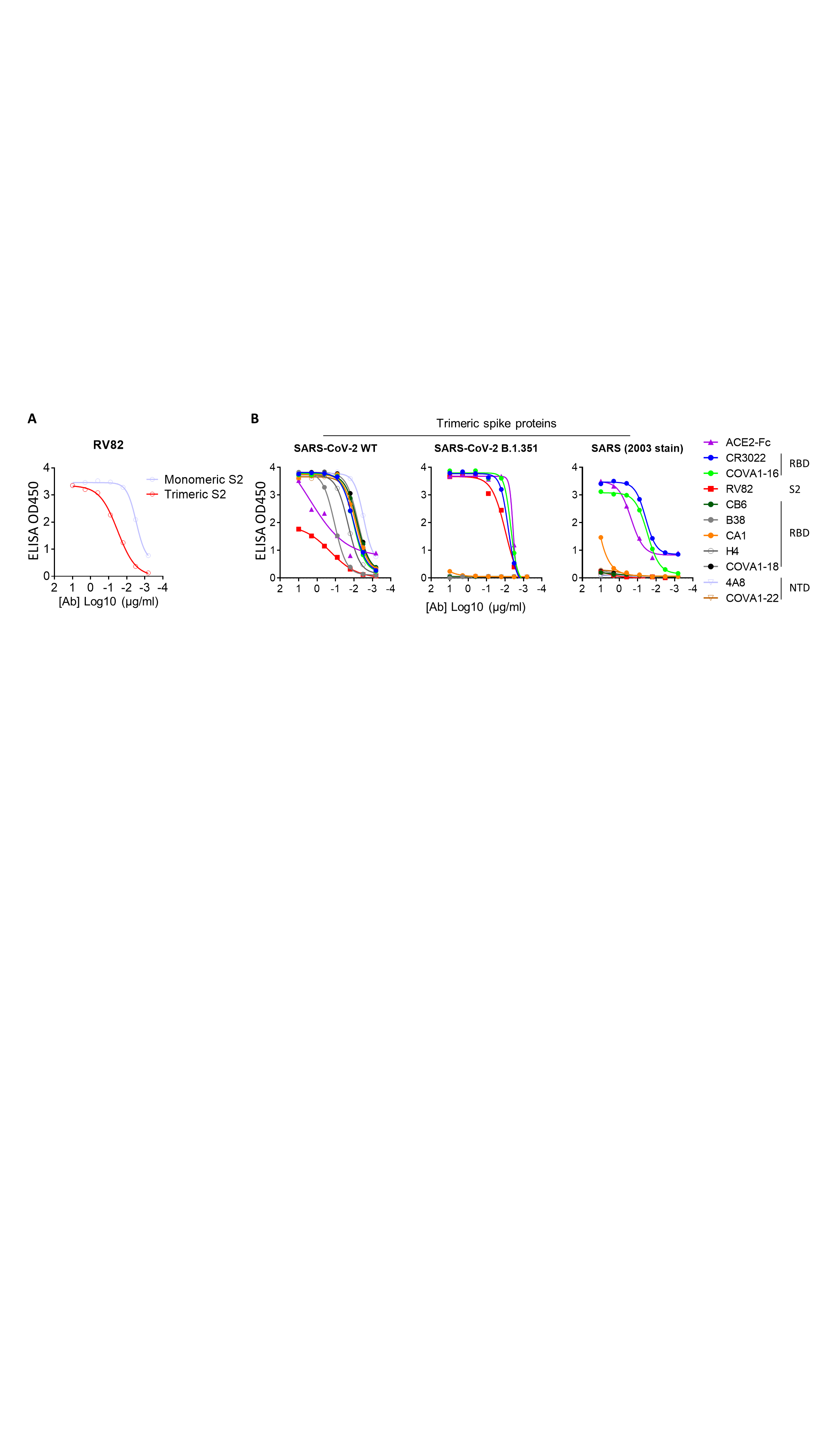
